# Ozone nanobubble treatments improve survivability of Nile tilapia (*Oreochromis niloticus*) challenged with a pathogenic multidrug-resistant *Aeromonas hydrophila*

**DOI:** 10.1101/2021.03.14.435289

**Authors:** Le Thanh Dien, Nguyen Vu Linh, Pattiya Sangpo, Saengchan Senapin, Sophie St-Hilaire, Channarong Rodkhum, Ha Thanh Dong

**Author notes:** Corresponding authors: H.T.Dong, C.Rodkhum.

## Abstract

Multidrug-resistant (MDR) bacteria has rapidly increased in aquaculture, which highlights the risk of production losses due to diseases and potential public health concerns. Previously, we reported that ozone nanobubbles (NB-O_3_) were effective at reducing concentrations of pathogenic bacteria in water and modulating fish immunity against pathogens; however, multiple treatments with direct NB-O_3_ exposures caused alterations to the gills of exposed-fish. Here, we set up a modified recirculation system (MRS) assembled with an NB-O_3_ device (MRS-NB-O_3_) to investigate whether MRS-NB-O_3_ were 1) safe for tilapia (*Oreochromis niloticus*), 2) effective at reducing bacterial load in rearing water, and 3) improved survivability of Nile tilapia following an immersion challenge with a lethal dose of MDR *Aeromonas hydrophila*. The results indicated no behavioral abnormalities or mortality of Nile tilapia during the 14 day study using the MRS-NB-O_3_ system. In the immersion challenge, although high bacterial concentration (~2 × 10^7^ CFU/mL) was used, multiple NB-O_3_ treatments in the first two days reduced the bacteria between 15.9% to 35.6% of bacterial load in water while bacterial concentration increased 13.1% to 27.9% in the untreated control. There was slight up-regulation of non-specific immune-related genes in the gills of the fish receiving NB-O_3_ treatments. Most importantly, this treatment significantly improved survivability of Nile tilapia with relative percent survival (RPS) of 64.7 - 66.7% in treated fish and surviving fish developed specific antibody against MDR *A. hydrophila*. In summary, the result suggests that NB-O_3_ is a promising alternative to antibiotics to control bacterial diseases, including MDR bacteria, and has high potential for application in recirculation aquaculture system (RAS).

**Highlights:**

- Multiple treatments of NB-O_3_ in a modified recirculation system (MRS) were relatively safe for juvenile Nile tilapia
- NB-O_3_ treatments in MRS significantly improved survivability of Nile tilapia challenged with multidrug-resistant (MDR) *A. hydrophila* with RPS of 64.7 - 66.7%
- Concentration of MDR *A. hydrophila* in MRS was reduced by 15.9 to 35.6% following each NB-O_3_ treatment, and increased by 13.1 to 27.9 % in untreated control
- Surviving fish developed specific antibody IgM against MDR *A. hydrophila*
- NB-O_3_ is a promising non-antibiotic approach to control diseases caused by MDR *A. hydrophila*

## 1. Introduction

Motile *Aeromonas* septicemia (MAS) is one of the most important bacterial diseases responsible for the loss of millions of dollars in the global freshwater aquaculture industry (da Silva et al., 2012; Hossain et al., 2014; Peterman and Posadas, 2019; Pridgeon and Klesius, 2012). The control of bacterial diseases still depends heavily on antibiotics. In recent years, a global issue of concern is the increase in antimicrobial resistant (AMR) bacteria as the consequence of misuse of antibiotics (Cabello, 2006; Cantas and Suer, 2014; Malik and Bhattacharyya, 2019). The high levels of AMR in the aquatic environment and aquaculture products pose a negative impact to not only aquaculture production, but also public health and international trade, especially in low- and middle-income countries (LMICs) where aquaculture is highly concentrated (Ben et al., 2019; Heuer et al., 2009; Okocha et al., 2018; Reverter et al., 2020). Currently, there is a high proportion of pathogenic multidrug-resistant (MDR) bacteria strains causing diseases in aquaculture (Santos and Ramos, 2018). In the battle to combat AMR, apart from alternatives to antibiotics, there are efforts to explore novel approaches for reducing the risk of bacterial diseases in aquaculture systems e.g. bacteriophage and nanobubble technology.

Nanobubbles (NBs) are bubbles less than 200 nm in diameter filled with chosen gases, neutral buoyancy, and having long residence time in the liquid solutions (Agarwal et al., 2011; Tsuge, 2014). Oxygen nanobubbles (NB-O_2_) have been used for improving dissolved oxygen (DO) in aquaculture systems, and promoting growth of Nile tilapia (*O. niloticus*) (Mahasri et al., 2018) and whiteleg shrimp (*Penaeus vannamei*) (Mauladani et al., 2020; Rahmawati et al., 2020). Recently, several studies have revealed that ozone nanobubbles (NB-O_3_) show promise at reducing quantities of pathogenic bacteria and improving DO in water, as well as modulating the immune systems against bacterial infections (Imaizumi et al., 2018; Jhunkeaw et al., 2021; Linh et al., 2021; Nghia et al., 2021).

Ozone is a powerful disinfectant that has been used to reduce concentrations of pathogens and improve water quality in both flow-through and recirculating aquaculture systems for many years (Powell and Scolding, 2018). However, low ozone solubility and poor stability are major reasons for low utilization efficiency. In addition, misuse of direct ozonation can critically impact aquatic organisms, resulting in behavioral abnormalities, changes in physiology, tissue damage, and mortality (Powell and Scolding, 2018). However, NBs technology has been reported to improve gas dissolvability in water and promote rapid oxidation of organic substances (Gurung et al., 2016). Hence, NB-O_3_ may enhance the solubility, stability, and efficacy of ozone in aquaculture systems (Fan et al., 2020). Kurita et al. (2017) reported that NB-O_3_ treatment significantly reduced planktonic crustacean parasites (63%) in juvenile sea cucumbers (*Apostichopus japonicas*) and sea urchins (*Strongylocentrotus intermedius*). In another study, NB-O_3_ demonstrated good disinfection of *Vibrio parahaemolyticus* in water, and prevention of acute hepatopancreatic necrosis disease (AHPND) in whiteleg shrimp (Imaizumi et al., 2018). We found that NB-O_3_ treatment (1-2 × 10^7^ bubbles/mL) reduced the level of *Streptococcus agalactiae* and *Aeromonas veronii* in water by more than 97% and made it relatively safe for juvenile Nile tilapia (Jhunkeaw et al., 2021). Most recently, we also reported that NB-O_3_ treatment modulated the innate immune defense system of Nile tilapia, and that pre-treatment of NB-O_3_ improved survivability of fish challenged with *S. agalactiae* (relative percent of survival of 60 −70%) (Linh et al., 2021). This finding suggests that NB-O_3_ may be a promising non-antibiotic treatment to control pathogenic MDR bacteria in aquaculture.

The limitations of direct application of NB-O_3_ with high level of ozone (3.5 mg/L, 970 mV ORP (oxidation reduction potential) is the tissue damage that this gas can cause to animals. Toxicity resulting in mortalities were reported for experimental shrimp in a study by Imaizumi et al. (2018). In our previous study on tilapia, we did not observe fish mortality but the fish gill morphology was damaged when fish were exposed directly to multiple NB-O_3_ treatments with an ORP range between 860 ± 42 and 885 ± 15 mV (Jhunkeaw et al., 2021). In this study, we set up a modified recirculation system coupled with ozone nanobubbles (MRS-NB-O_3_). Subsequently, we evaluated the system to determine if it was effective at suppressing pathogenic MDR *A. hydrophila* and the survivability of juvenile Nile tilapia.

## 2. Materials and methods

### 2.1. Bacterial strains and culture conditions

A laboratory strain of multidrug resistant *A. hydrophila* BT14, isolated from an outbreak of MAS in 2018, was used in this study. Briefly, this bacterial strain was identified by Matrix-Assisted Laser Desorption/Ionization-Time of Flight Mass Spectrometry (MALDI-TOF MS) and PCR-sequencing using *gyrB* housekeeping gene, following previous studies (Anand et al., 2016; Navarro and Martínez-Murcia, 2018). Based on the method proposed by Magiorakos et al. (2012), *A. hydrophila* BT14 was identified as a multidrug-resistant bacterium due to the fact that it resisted at least three classes of antimicrobials, including Ampicillin 10 μg (Penicillins), Tetracycline 30 μg (Tetracyclines), and Sulfamethoxazole-Trimethoprim 23.75 −1.25 μg (Folate pathway inhibitors) (Table S1). For the bacterial challenge test, MDR *A. hydrophila* BT14 was propagated in 1 L of TSB at 28 °C with 18 h shaking-culture at 150 rpm. The bacterial concentration was determined by conventional plate count method (Harrigan and McCance, 2014).

### 2.2. Experimental fish

Healthy Nile tilapia (3.92 ± 1.01 g) from a commercial tilapia hatchery in Thailand were acclimated in dechlorinated tap water for 2 weeks at 29 ± 1.0 °C before the experiments. Fish were fed with commercial tilapia feed (crude-protein 30%) at rate of about 3% of fish weight twice daily. Before starting the experiments, ten fish were randomly selected for bacterial isolation and found to be free of *A. hydrophila*. The experiments on animals were conducted with permission of Thai Institutional Animal Care and Use Committee (Approval no. MUSC62-039-503).

### 2.3. MRS-NB-O_3_ system setup and water parameter measurement

The ozone nanobubble system consisted of an oxygen concentrator (Model: VH5-B, Shenyang Canta Medical Technology Company Limited, Liaoning, China) connected to an ozone generator (Model: CCba15D, Coco Technology Company Limited, Chonburi, Thailand) and a nanobubble generator (Model: aQua+075MO, AquaPro Solutions Private Limited Company, Singapore). The NB-O_3_ system was attached to a modified recirculation system (MRS) which contained two 100 L-fiberglass tanks (50 L dechlorinated tap water in each tank) that exchanged water by water pumps. One tank received the NB-O_3_, the other tank housed the fish (Figure 1). All water quality parameters were measured in triplicate in the MRS-NB-O_3_. Water temperature, pH, dissolved oxygen (DO), and oxidation reduction potential (ORP) were measured and compared from both tanks using a multi-parameter meter (YSI Professional Plus, YSI Incorporated, USA). During the application of the NBs, water samples were collected at 0 min, 5 min, 10 min of NB-O_3_ treatment and 30 min post-treatment for measurement of dissolved ozone (ppm-mg/L) using K-7434 Ozone Vacu-vials Kit (Oxidation Technologies, USA).

**Figure 1.**
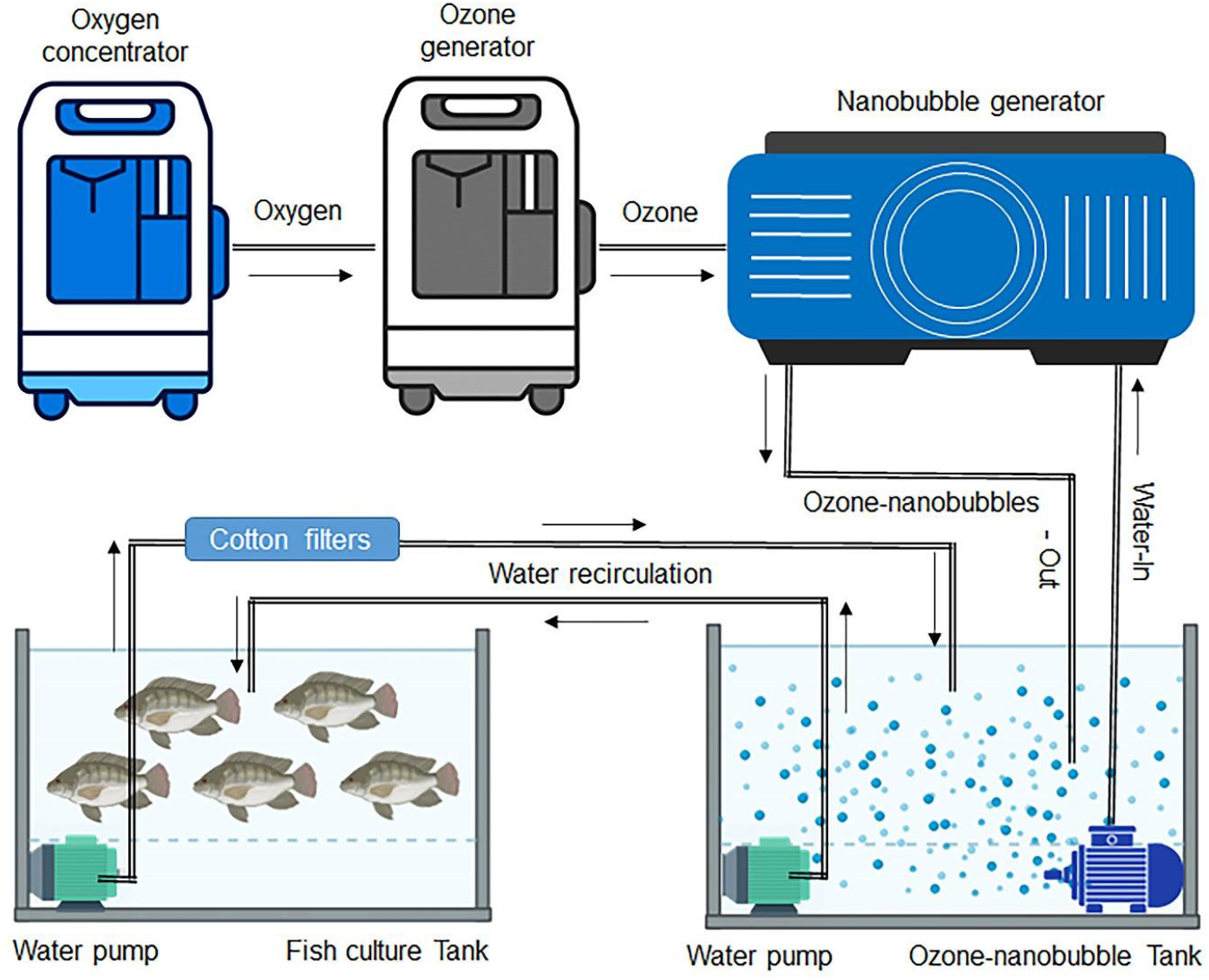
Experimental set-up of MRS-NB-O_3_. Oxygen concentrator releases oxygen as a material to synthesize ozone using ozone generator. Ozone was lead to nanobubble generator. Inside the system, ozone was diffused in nanobubble water and released to ozone-nanobubble tank. Thereafter, NB-O_3_ water was pumped to fish culture tank. The rearing water were recirculated between NB-O_3_ tank and fish culture tank via a pump system assembled to cotton filter box to absorb fish feces and leftover feed.

### 2.4. Effect of MRS-NB-O_3_ on fish safety

To evaluate the safety of Nile tilapia juveniles cultured in MRS-NB-O_3_ system, 136 fish were divided into four tanks (50L dechlorinated tap water per tank) consisting of two replicate groups (controls and MRS-NB-O_3_) with 34 fish per tank. The treatment group was treated with NB-O_3_ (oxygen input 2 L per min) 7 times (10 min/time) at 1, 12, 24, 36, 48, 60, and 72 h from the start time of the experiment. Aeration was provided one hour after each treatment. The control group was treated with normal aeration instead of NB-O_3_. Fish were observed every 12 h for behavioral abnormality and mortality over a 14-day period. The water parameters including temperature, pH, DO, and ORP were measured before and during treatment. After every treatment, two fish in each tank were randomly collected and preserved for gill histology examination. Formalin preserved samples (n = 28) were subjected to routine histology. The histopathological changes were observed under the Leica DM1000 digital microscope equipped with a digital camera DFC450 (Leica, Singapore).

### 2.5. Immersion challenge trial for MDR *A. hydrophila* BT14

To establish the immersion challenge dose, 80 fish were divided into four 50 L tanks, each tank containing 20 fish. Three tanks were challenged with MDR *A. hydrophila* BT14 by adding 1 L of bacterial culture (approx. 8 × 10^6^, 8 × 10^7^, and 8 × 10^8^ CFU/mL) to each tank to reach the final concentrations of 2 × 10^5^, 2 × 10^6^, and 2 × 10^7^ CFU/mL, respectively. A total 1 L of culture medium without bacteria was added to a negative control tank. Air-stones were used in all tanks for air supply and approximate 50% of the water was changed after 48 h. Clinical signs of MAS and mortalities were recorded every 12 h for 14 days. The representative dead or moribund fish were subjected to bacterial re-isolation using selective medium Rimler Shotts (RS, Himedia, India) supplemented with Novobiocin (Oxoid, UK).

### 2.6. Effect of multiple NB-O_3_ treatments in MRS on Nile tilapia challenged with MDR *A. hydrophila*

#### Fish survivability, gill collection, and water collection

Two trials were conducted to test the effect of our MRS NB-O_3_ treatments. In the first trial, 128 fish were randomly divided into four groups (32 fish per tank): Group 1 was exposed to culture medium without NB-O_3_ treatment (no Ah + no NB-O_3_); Group 2 was exposed to bacteria without NB-O_3_ (Ah + no NB-O_3_); Group 3 was exposed to culture media only and treated with NB-O_3_ (no Ah + NB-O_3_); Group 4 was challenged with *A. hydrophila* and treated with NB-O_3_ (Ah + NB-O_3_).

In bacterial challenge groups 2 and 4, 1 L of MDR *A. hydrophila* BT14 (approx. 8 × 10^8^ CFU/mL) was added to 50 L water to reach a final concentration of approx. 2 × 10^7^ CFU/mL. The fish were maintained at 29 ± 1 °C with aeration for 3 h. Afterwards, fish in groups 3 and 4 were treated for 10 min with NB-O_3_ at 1, 12, 24, 36, and 48 h post-challenge, while group 1 and group 2 were treated with normal aeration. In order to investigate the effect of NB-O_3_ treatments on the fish immune response in our MRS, the gills from 4 fish were randomly sampled at 3 h after the 1^st^, 2^nd^, and 3^rd^ NB-O_3_ treatments and preserved in 200 μL of Trizol reagent (Invitrogen, USA) for immune genes analysis. The remaining fish were observed daily for 14 days and mortality was recorded. Representative moribund or freshly dead fish were collected for bacterial re-isolation using Rimler Shotts (RS) medium plus Novobiocin as described above. The relative percent survival (RPS) was calculated according to the formula described by Ellis (1988): RPS = [1 −(% mortality in challenge/ % mortality in control)] × 100. In parallel, water samples from groups 2 and 4 (challenged with *A. hydrophila*) were evaluated for bacterial enumeration using conventional plate count method (Harrigan and McCance, 2014). The percentage of bacterial fluctuation was calculated based on bacterial concentration (CFU/mL) before and after NB-O_3_ treatment.

In the second trial, the experiment was repeated in the same manner as the first with the exception that 20 fish were used for each group and this experiment focused mainly on monitoring survival rate and bacterial enumeration. This experiment was repeated to confirm our initial survival results in the first trial.

#### Visualization of live and dead bacteria before and after treatment with NB-O_3_

A volume of 25 mL water in group 4 (Ah + NB-O_3_) was sampled before and after the first NB-O_3_ treatment for assessment of the viability of *A. hydrophila*. A bacterial suspension was prepared and stained following the protocol of LIVE/DEAD *Bac*light Bacterial Viability Kit (Cat. No. L7012, Thermo-Fisher Scientific, USA). In brief, the bacterial suspension were centrifuged at 10,000 x *g* for 10 min at 4°C. The pellets were collected and re-suspended in 2 mL of sterile normal saline buffer, incubated at room temperature for 1 h, mixing every 15 min. Bacteria were washed two times by centrifugation at 10,000 x *g* for 10 min at 4°C and pellet resuspension was done in 20 mL and 10 mL of sterile normal saline buffer for the first and second time of washing. Staining processes were conducted by mixing 1.5 μL of SYTO^®^9, 1.5 μL of Propidium Iodine (PI), and 1 mL of bacterial suspension in a microtube. The mixture was incubated at room temperature in the dark for 15 min. After that, 5 μL mixtures were pipetted onto glass slides, covered with a coverslip and examined under a confocal laser scanning microscope CLSM (Model: DM1000, Leica Microsystem Private Limited Company, Singapore) assembled with incident light fluorescence to visualize live and dead bacteria. Five random fields from each slide were imaged. Fluorescence signals were counted in ImageJ software based-on Watershed algorithm.

#### Expressions of innate immune-related genes

To investigate expression of innate immune-related genes, total RNA of gill samples was extracted using Trizol reagent (Invitrogen, USA) following the manufacturer’s instructions. The first complementary DNA (cDNA) strand was synthesized from 2.0 μg of the total RNA using iScript^™^ Reverse Transcription Supermix (Bio-Rad, USA) according to the procedure described in the product manual. Quantitative real-time PCR (qPCR) using SYBR green reagent (iTaq^™^ Universal SYBR^™^ green Supermix, Bio-Rad, Hercules, CA, USA) was carried out using primers specific for 3 immune genes (Table 1). The qPCR amplification cycles were performed using a CFX Connect^™^ Real-time System (Bio-Rad, USA). Cycling conditions were 94 °C for 15 s, 40 cycles of denaturation at 95 °C for 30 s, annealing at the optimal temperature of each primer as indicated for 30 s, and a final extension at 72 °C for 30 s. Melting curves were obtained in the 55 to 85°C range with 0.1 °C increments per second to evaluate for the specificity of all qPCR products. The qPCR data will be analyzed using the 2^-△△Cq^ method (Livak and Schmittgen, 2001). The transcript levels of each target gene were obtained as C_q_ values and normalized to that the *EF-1a* as an internal reference.

**Table 1.**
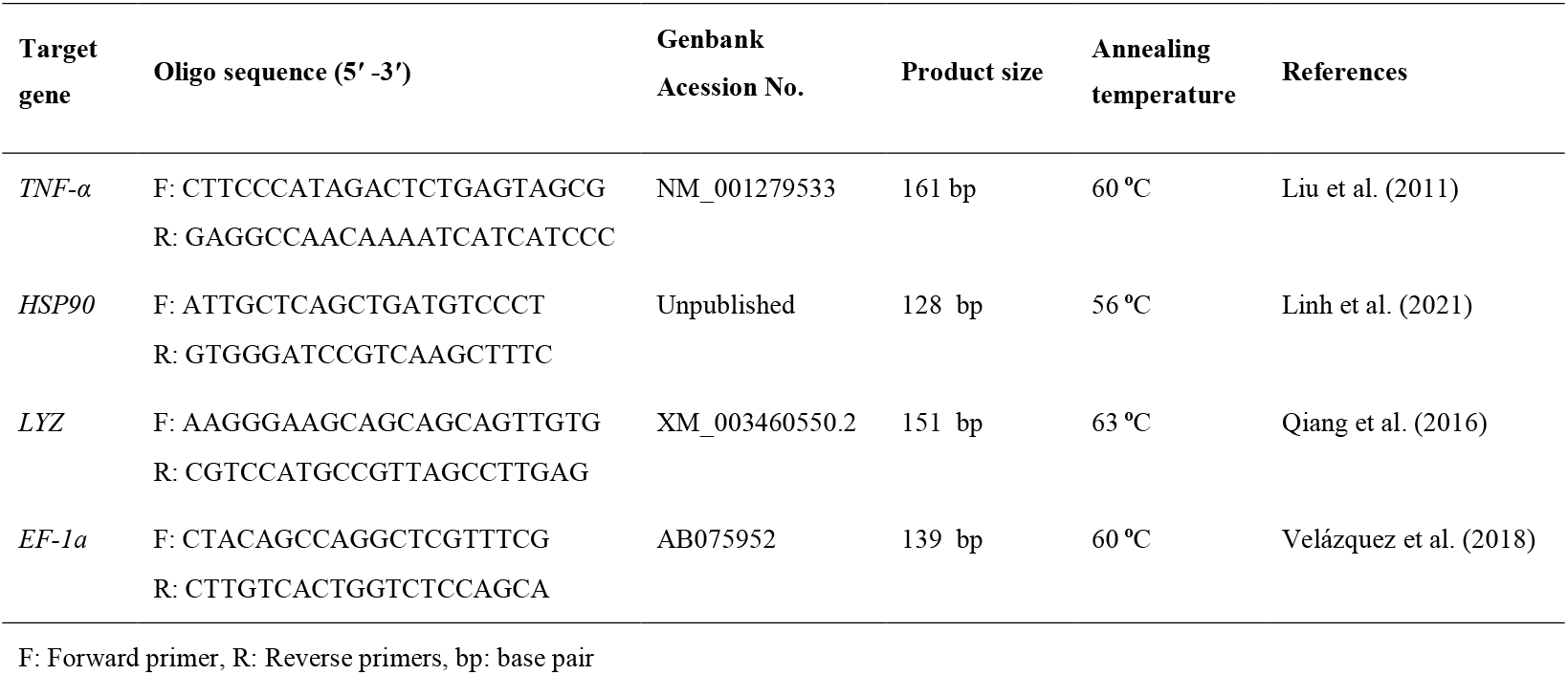
Primers used to quantify relative gene expression in this study

#### Determination of serum antibody by the enzyme-linked immunosorbent assay (ELISA)

In order to determine whether surviving fish at day 14 post challenge develop specific antibodies (IgM) against *A. hydrophila*, blood samples were collected from fish in the first trial (four from Ah + no NB-O_3_ group and five from each of the other groups). Blood samples were kept at room temperature for 1 h before being centrifuged at 8.000 x g for 15 min. The collected fish sera were stored at −20°C until used. An ELISA was carried out following the protocol described by Linh et al. (2021) with minor modification. In brief, 96 well EIA/RIA plates (Costar^®^, Corning Inc., USA) were coated with formalin-killed *A. hydrophila* whole-cell antigen (OD_600nm_ = 1.0). Fish sera (dilution 1:256), anti-Tilapia IgM secondary antibody (1:200) (Soonthonsrima et al., 2019), and commercial goat anti mouse antibody horseradish peroxidase (HRP) conjugate (1:3000) were used for the ELISA assay in this study and samples were read at an absorbance of 450 nm using a SpectraMax^®^ iD5 Multi-Mode Microplate Reader (Molecular Devices, USA).

### 2.7. Statistical analysis

Cumulative mortality and percent survival data from the challenge experiments were analyzed by the Kaplan-Meier method and differences among groups were tested using a log-rank test, *p*-values of 0.05 or less were considered statistically significant. Fish innate immune-related gene expression was analyzed by ANOVA, *p*-values of 0.05 or less were considered statistically significant. Duncan’s post-hoc test was used to measure specific differences between pairs of mean. The OD_450nm_ readings from our indirect ELISA assay were analyzed using a Kruskal-Wallis test, *p*-values of 0.05 or less were considered statistically significant. Multiple comparison analyses were performed by Bonferroni test. All statistical analyses were performed using SPSS Software ver22.0 (IBM Corp., USA).

## 3. Results

### 3.1. Effect of MRS-NB-O_3_ on water parameters

For the 10 min NB-O_3_ treatment in the MRS, the change of water parameters, including temperature, pH, DO, and ORP, are displayed in Figure 2. Temperature and pH values appeared stable over time in both the NB-O_3_ treated tank and the culture tank (which did not have fish for this investigation). The DO increased significantly after 10 min NB-O_3_ treatments in both tanks. The DO level in the culture tank increased from 5.07 ± 1.61 to 13.97 ± 0.84 mg/L (increase of 8.9 mg/L), while there was an higher increase in NB-O_3_ tank (from 6.84 ± 1.08 to 19.74 ± 1.28 mg/L). The significantly different trend of ORP value was observed in the NB-O_3_ treated tank and culture tank. The ORP decreased slightly from 424.9 ± 24 to 396 ± 61.9 mV in fish culture tank, whereas the ORP in NB-O_3_ tank increased rapidly from 417.7 ± 23.6 to 791.7 ± 71.5 mV after 5 min NB-O_3_ treatment and reached 870.1 ± 12.4 mV after 10 min. During NB-O_3_ treatment, dissolved ozone concentration at 0 min, 5 min, and 10 min in treated tank were 0.02, 1.16, and 1.37 mg/L respectively, whereas significantly lower values, 0.03, 0.06, and 0.14 mg/L were recorded in system’s fish culture tank at the same time points. At 30 min post-treatment, dissolved ozone concentration in NB-O_3_ treated and fish culture tanks decreased to 0.05 and 0.03 mg/L respectively.

**Figure 2.**
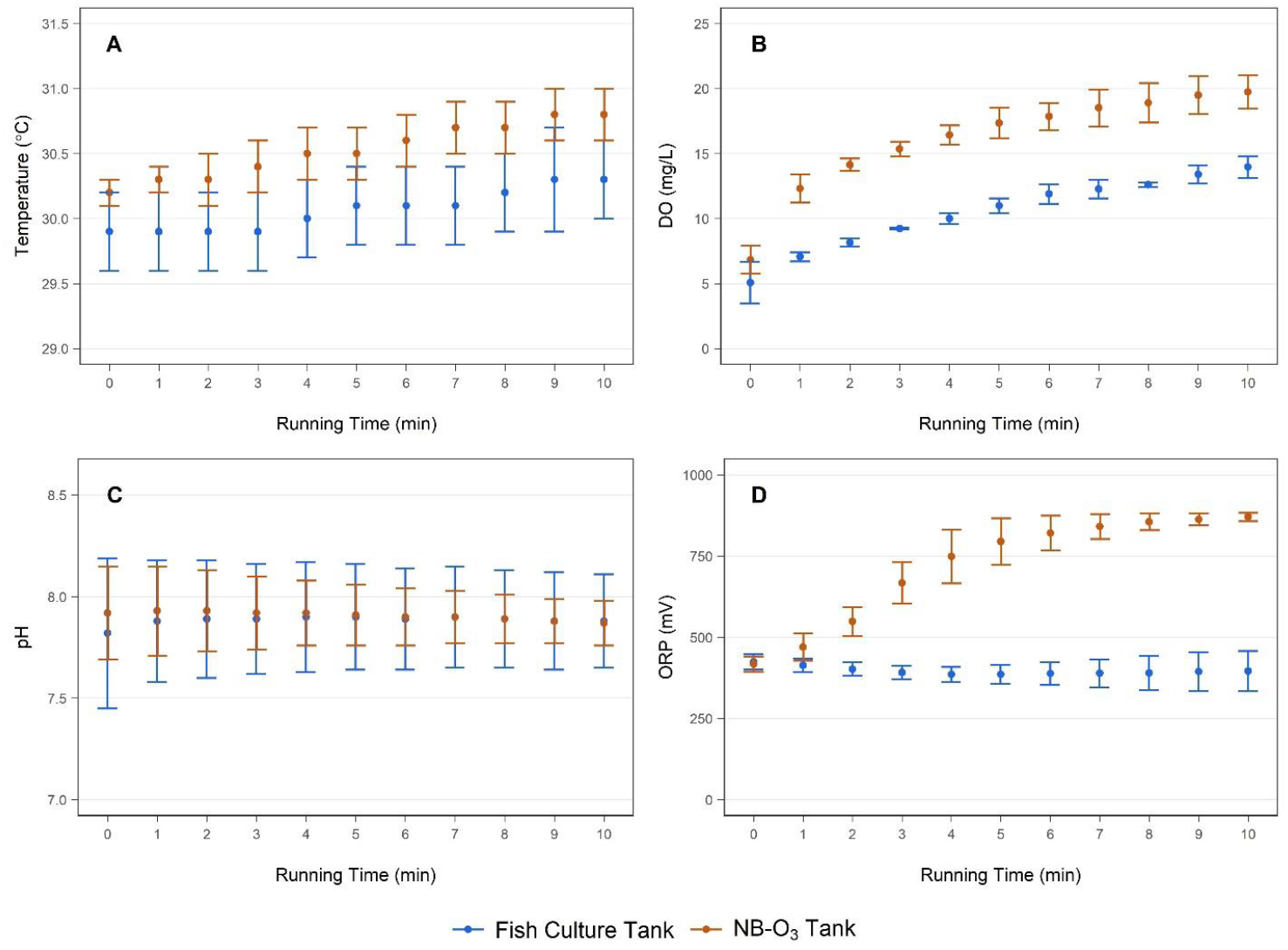
Measurement of water parameters including temperature (**A**), DO (**B**), pH (**C**), and ORP (**D**) during 10 min NB-O_3_ treatment with 2 L/min oxygen input in MRS. Value of water parameters are mean ± SD (n = 3).

### 3.2. Effect of MRS-NB-O_3_ on fish safety

No mortality or behavioral abnormalities in fish were observed in either the control and NB-O_3_ treated groups during and after treatments. All fish survived the 14 day study period. Histologically, there were no differences in gill morphology in control and treatment groups after five NB-O_3_ treatments. However, alterations were observed in the gill filaments after the 6^th^ and 7^th^ treatments (Figure S1). The fluctuation of water parameters was consistently similar during every treatment (Table S2), and similar to the trend in the previous experiment without fish (Figure 2). Temperature and pH increased slightly in both groups during treatment. Dissolved oxygen in the fish culture tanks of the MRS-NB-O_3_ increased significantly from 4.98 −6.97 mg/L (before each treatment) to 12.26 −15.33 mg/L (at each 10 min of treatment) and dropped to 9.28 −12.69 mg/L after the 10 min treatment. ORP values in fish culture tanks did not increase and remained relatively stable in control and NB-O_3_ treated groups.

### 3.3. Immersion challenge trial for MDR *A. hydrophila* BT14

The cumulative mortality of Nile tilapia challenged with three different doses of MDR *A. hydrophila* BT14 by immersion was dose-dependent (Figure 3). The fish challenged with 2 × 10^7^ CFU/mL (high dose) had a 75% mortality rate, and death occurred mainly in the first 4 days of the experiment. In the 10-fold lower dose, there was only 25% mortality and most fish died from days 4 to 9. There was no mortality in the group challenged with 2 × 10^5^ CFU/mL or the control group (Figure 3). The clinically sick fish showed lethargy, loss of appetite, and tended to swim at the surface, but did not reveal significant external or internal symptoms except pale livers. Bacterial isolation from representative dead fish (n = 5) revealed dominant colonies of bacteria, morphologically resembling *A. hydrophia* on selective medium. From this result, the dose of 2 × 10^7^ CFU/mL was used for subsequent challenge assays.

**Figure 3.**
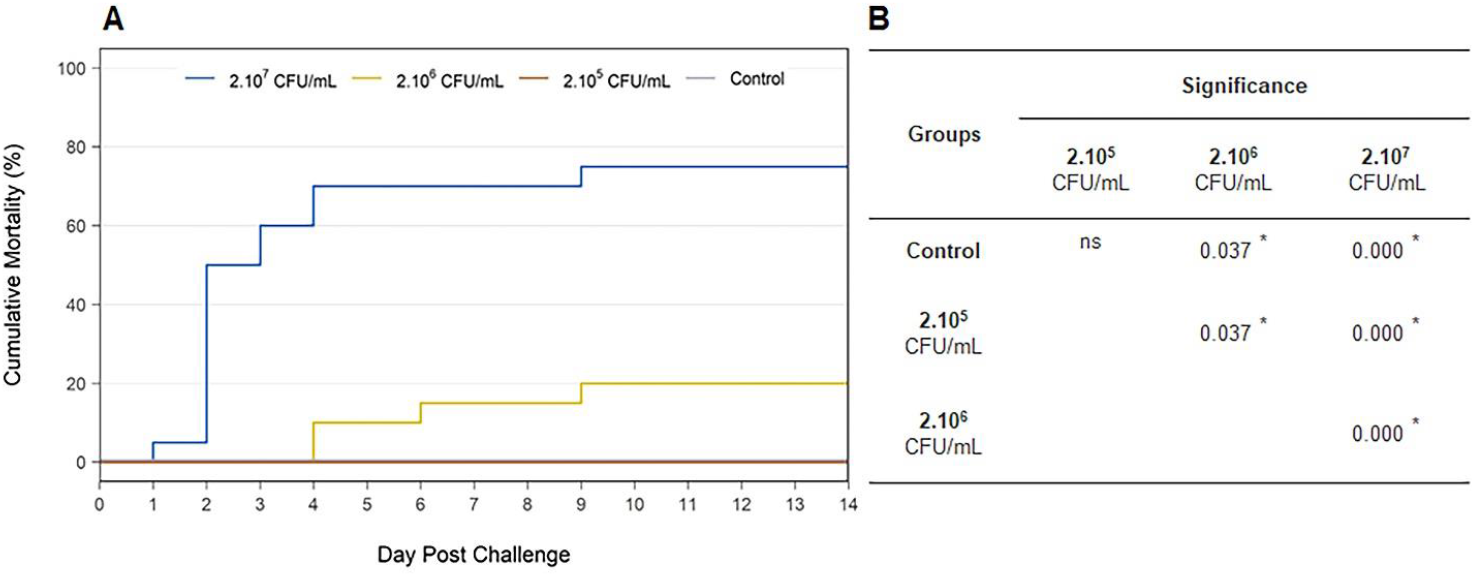
Kaplan-Meier analysis of cumulative mortality of Nile tilapia (n = 20) challenged with 3 doses of MDR *A. hydrophila* BT14 by immersion method (A). Control was exposed to culture medium without bacteria. Differences between groups were tested using log-rank test shown in (B). “*” denotes significant difference (*p* < 0.05), “ns” means not significant.

### 3.4. MRS-NB-O_3_ improved survivability of Nile tilapia challenged with the MDR *A. hydrophila* BT14

The results of the challenge tests were consistent between replicates (Figure 4). The group challenged with *A. hydrophila* followed by NB-O_3_ treatments (Ah + NB-O_3_) had 70 and 75% survival compared to 15 and 25% in the group challenged with bacteria receiving no NB-O_3_ treatment (Ah + no NB-O_3_). This difference was statistically significant (*p* = 0.001) in both trials. No mortality was observed in the negative control group (no Ah + no NB-O_3_) during the 14 day study period. However, there were 5 and 15 % mortality in the control groups treated with NB-O_3_ without a precedent bacterial challenge (no Ah + NB-O_3_). However this was not statistically significant to the negative control group in either trials (*p* = 0.317 in trial 1 and *p* = 0.075 in trial 2 (Figure 4)). The relative percent survival (RPS) of NB-O_3_ treatments in the 2 replicate treatment groups were 64.7 and 66.7%, respectively.

**Figure 4.**
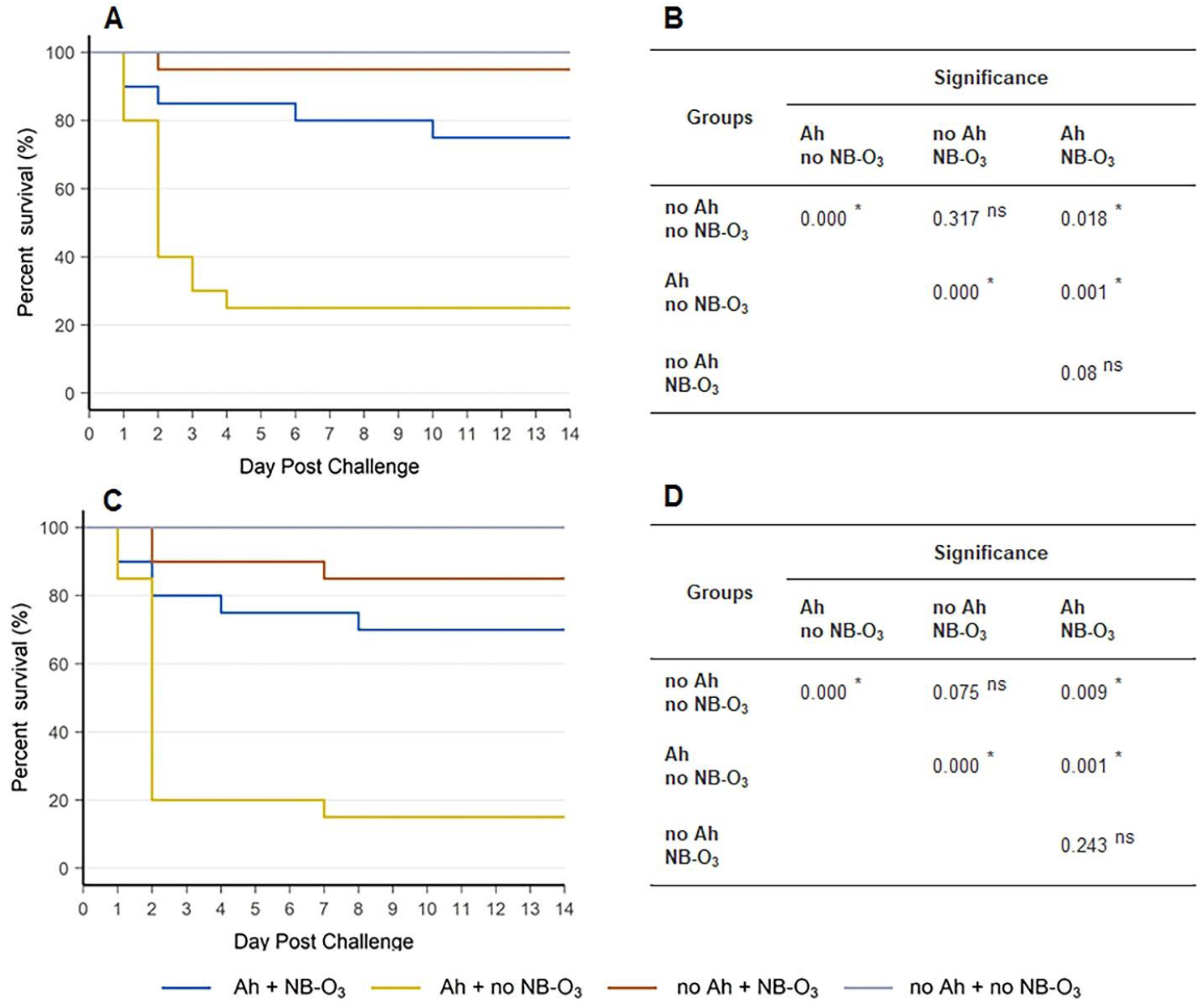
Kaplan-Meier analysis of percentage survival of Nile tilapia (n = 20) challenged with MDR *A. hydrophila* BT14. The experiment was done in two independent trials, trial 1 (A) and trial 2 (C). Differences between groups in each trial were tested using log-rank test shown in (B) and (D) respectively. “*” denotes significant difference (*p* < 0.05), “ns” means not significant.

The moribund fish in challenge groups showed pale liver and behavioral abnormalities, including lethargy, loss of appetite, and surface swimming. The typical colonies of *A. hydrophila* were consistently recovered from internal organs (i.e. liver, kidney) of representative dead fish using RS medium supplemented with Novobiocin.

In parallel, bacterial concentration in the water column was monitored in two groups challenged with *A. hydrophila*. In the group Ah + NB-O_3_, bacterial load in fish culture tanks after the 1^st^, 2^nd^ and 3^rd^ treatments were reduced by 35.6, 23.3, and 20.2%, respectively in the first trial, and by 23.9, 21.1, and 15.9%, respectively in the second trial (Figure 5). By contrast, bacterial load in the Ah + no NB-O_3_ increased by 13.4, 13.1, and 27.1% in the first trial, and by 15.6%, 19.8, and 27.9 % during the same time period in the second trial. Representative photomicrographs of comparative visualization of live and dead bacteria before and after treatment with NB-O_3_ are illustrated in Figure 6. Before NB-O_3_ treatment, the majority of bacterial cells appeared to be alive (i.e. stained fluorescent green), with very few dead cells (i.e. red color) (Figure 6A-C). However, after 10 min NB-O_3_ treatment, the density of dead cells (red staining cells) increased considerably (17.45%) per microscopic field.

**Figure 5.**
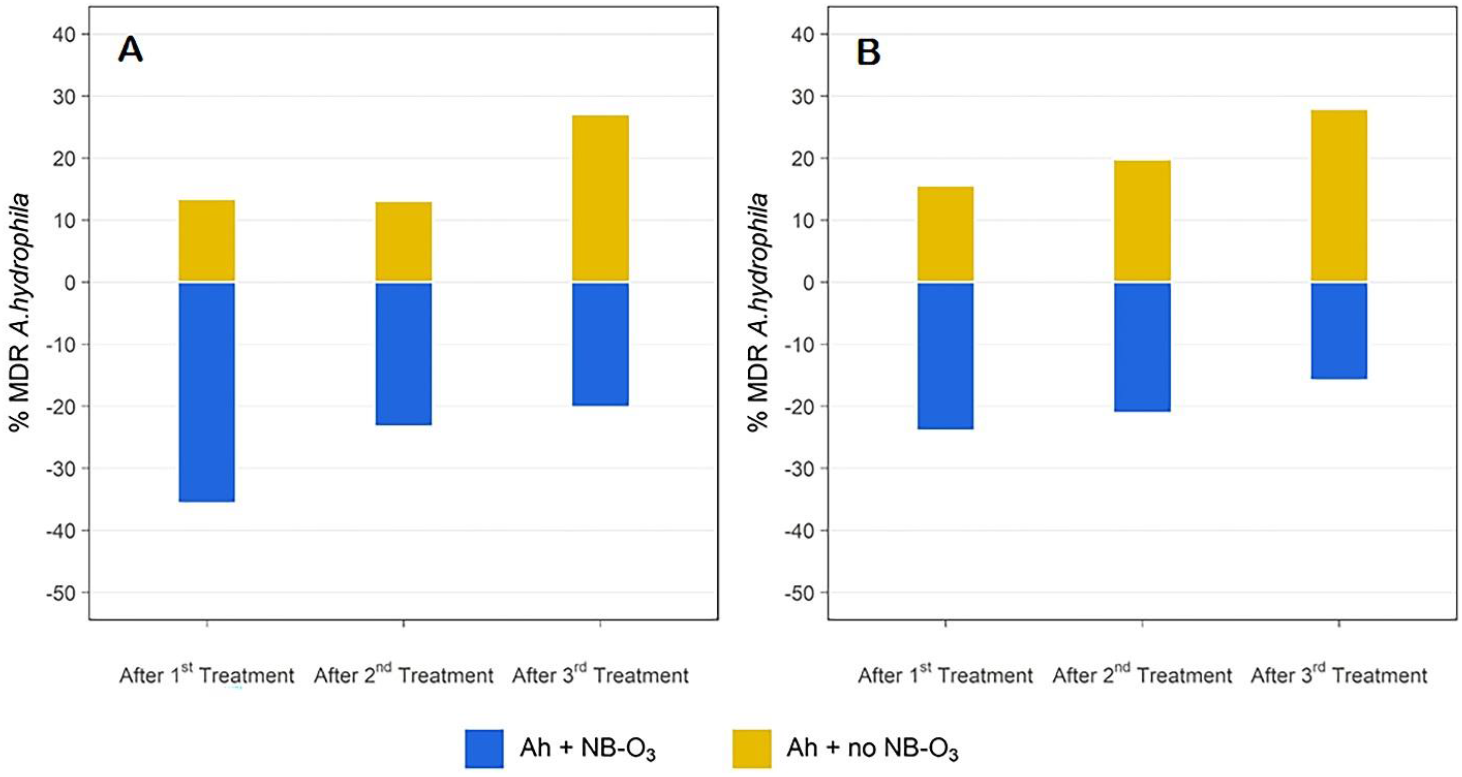
Concentration of MDR *A. hydrophila* BT14 in rearing water between un-treated and treated by 10 min NB-O_3_ groups after the 1^st^, 2^nd^, and 3^rd^ treatment. A, trial 1; B, trial 2.

**Figure 6.**
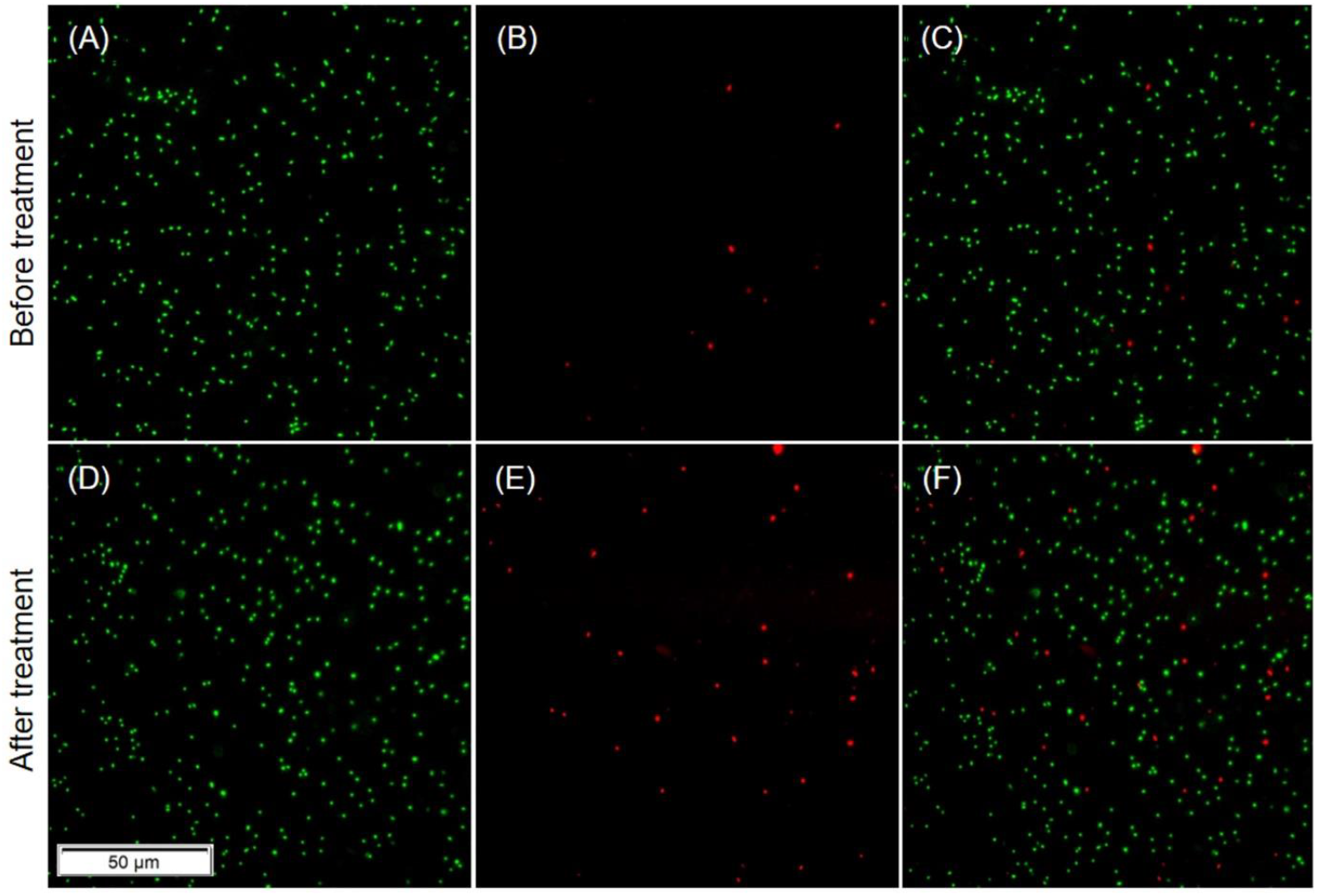
Confocal scanning laser microscope image of MDR *A. hydrophila* BT14 viability following the 1^st^ treatment with 10 min NB-O_3_ (A-C: before 1^st^ treatment and D-F: after 1^st^ treatment). Figure C is merged by A and B whereas figure F is merged by D and E. Green fluorescent indicates live bacterial cells and red fluorescent indicates dead bacterial cells using LIVE/DEAD *Baclight* Bacterial Viability Kit with two staining reagents SYTO^®^9 and PI.

### 3.5. Expressions of innate immune-related genes

The expression levels of innate immune genes from different groups after each NB-O_3_ treatment are shown in Figure 7. Although not statistically significant, the overall expression levels of immune genes *LYZ, HSP90*, and *TNF-a* in the gills of the fish exposed to NB-O_3_ treatments tended to be slightly higher than that of the untreated control, except for the first treatment. Specifically, the trends included *LYZ* expression in treated group with or without *A. hydrophila* challenge which rose after the 2^nd^ and 3^rd^ treatment compared to that in the negative control group. The highest expression level (approx. 2.2 folds) was recorded in NB-O_3_ treated group with *A. hydrophila* at the 3^rd^ treatment. Expression of *HSP90* had different patterns for different experiments. The expressions in NB-O_3_ treated group with or without *A. hydrophila* challenge increased at the 2^nd^ treatment but decreased similar to the levels in the control group for the 3^rd^ treatment. The relative transcription level of *TNF-a* increased slightly (1.4 fold) with the highest expression level in NB-O_3_ treated group.

**Figure 7.**
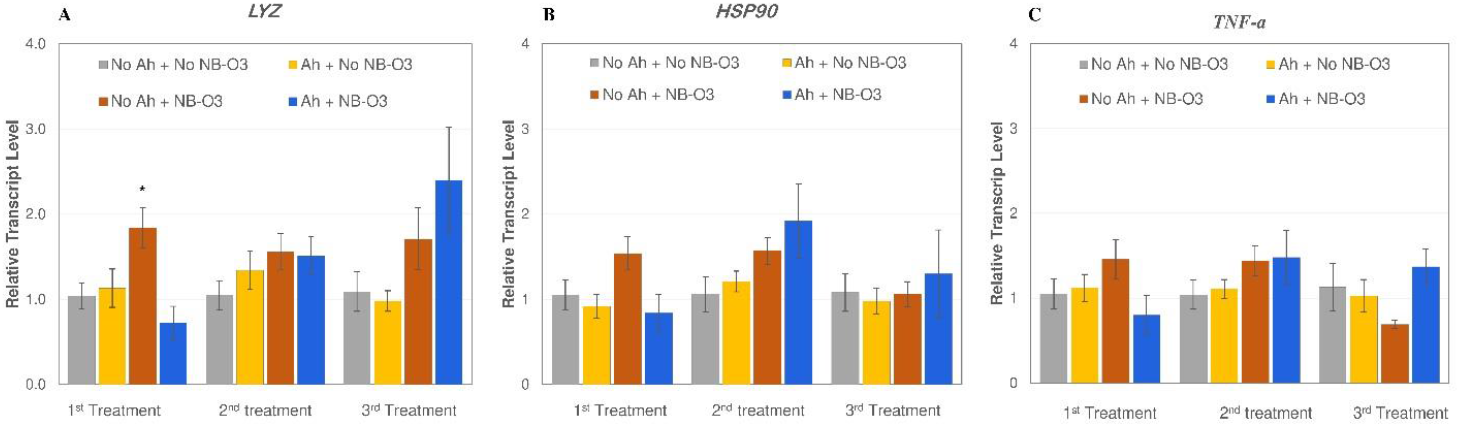
Relative expression of *LYZ* (**A**), *HSP90* (**B**) and *TNF-a* (**C**) in fish gills in 4 groups: no Ah + no NB-O_3_, Ah + no NB-O_3_, no Ah + NB-O_3_ and Ah + NB-O_3_ after 1^st^, 2^nd^ and 3^rd^ treatment with NB-O_3_. The expression of target genes was normalized using *EF-1a*. Value of relative transcript level are mean ± a standard error of the mean (SEM) bar (n = 4) and “*” above the bar indicates significant difference between groups (*p* < 0.05).

### 3.6. Specific antibody (IgM) response post-challenge

All surviving fish in both groups challenged with MDR *A. hydrophila* had significantly higher levels of specific antibody (IgM) compared to the two unchallenged control groups (*p* < 0.05) as measured by indirect ELISA (Kruskal-Wallis test: H (3) = 15.542, *p* = 0.001). The serum from fish in the Ah + NB-O_3_ group had the highest OD_450_ readings (0.44 ± 0.076), followed by OD readings of serum in Ah + no NB-O_3_ group (0.42 ± 0.06). In contrast, the lowest level (0.06 ± 0.004) was recorded in the negative control (no Ah + no NB-O_3_). A higher level but not statistically significant difference with negative control was shown in group no Ah + NB-O_3_ (0.1 ± 0.013) (Figure 8).

**Figure 8.**
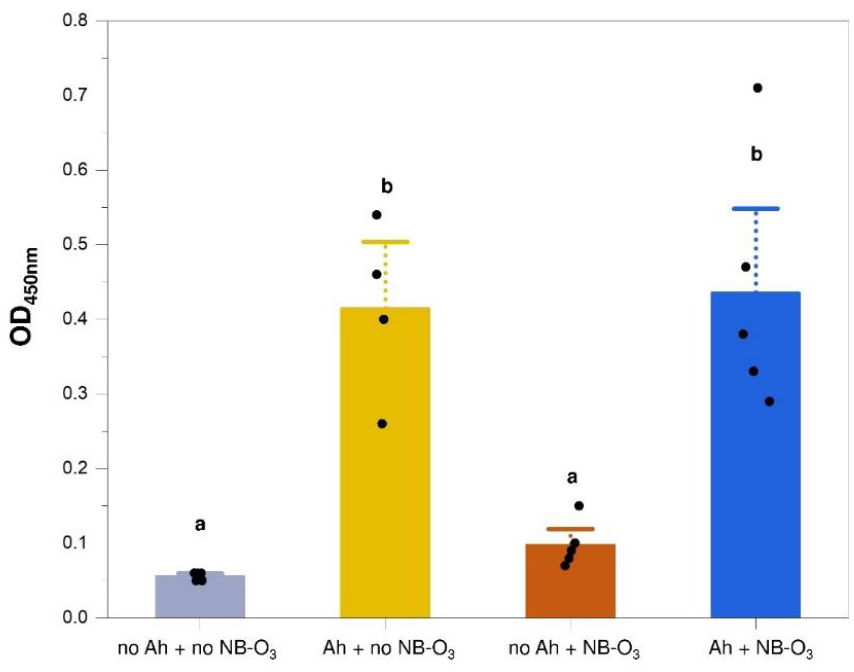
Indirect ELISA analysis of *A. hydrophila* specific IgM antibody. Fish sera were collected on day 14 and 1:256 dilutions were used to test for antigen specific IgM. Data were expressed as mean absorbance at OD_450nm_ with a SEM bar. One dot represents one biological replicate (n = 4 in group Ah + no NB-O_3_, n = 5 in other groups). Different letters above the bar indicate significant difference between groups (*p* < 0.05).

## 4. Disscussion

Several studies have reported potential applications of NB-O_3_ for pathogen disinfection in aquaculture water to reduce the risk of infectious diseases in both fish and shrimp (Imaizumi et al., 2018; Jhunkeaw et al., 2021; Kurita et al., 2017). We recently reported an additional benefit of NB-O_3_ in modulating of the innate immune defense system in Nile tilapia to fight against *S. agalactiae* (Linh et al., 2021). However, all the precedent studies exposed the animals directly to NB-O_3_ (NB-O_3_ was exposed directly into the tank containing fish or shrimp) and this resulted in mild to severe health impacts on the exposed animals. High dose of ozone (960 mV ORP) were toxic to shrimp (Imaizumi et al., 2018), or caused gills alteration in tilapia after repeated exposures to NB-O_3_ (~860 mV ORP) (Jhunkeaw et al., 2021). Therefore, we modify a NB-O_3_ system on a laboratory scale to better understand this technology and overcome this drawback.

Ozone is an unstable molecule, even in the form of nanobubbles, which degrades relatively quickly (Jhunkeaw et al., 2021). Based on this characteristic, we set up a modified recirculation system coupled with NB-O_3_ technology (MRS-NB-O_3_), which separated the NB-O_3_ treatment tank from the culture tank containing fish to reduce direct exposure of the fish to high level of ozone. Interestingly, during treatment, ozone level increased rapidly in the NB-O_3_ treatment tank but did not increase in the fish culture tank, as indicated by ORP values (870.1 ± 12.4 vs. 396 ± 61.9 mV ORP) and dissolved ozone concentrations (1.37 vs. 0.14 mg/L). Several studies suggested that ORP levels in the range from 300 to 425 mV ORP were safe for fish, crustaceans, and molluscs (Li et al., 2014; Powell and Scolding, 2018; Stiller et al., 2020). In the MRS-NB-O_3_ set up, multiple treatments (up to seven 10 min treatments) in this study appeared to be relatively safe for juvenile Nile tilapia, with no mortality over a 14 day period. We also noticed that the MRS-NB-O_3_ system could avoid excess DO level in the culture tank that commonly occurred when the NB-O_3_ treatments were applied directly to the fish tanks (Jhunkeaw et al., 2021).

This study revealed that multiple NB-O_3_ treatments in our MRS-NB-O_3_ system improved survivability of Nile tilapia (*O. niloticus*) challenged with a pathogenic multidrug-resistant *A. hydrophila*. Motile Aeromonads have been reported as one of the most common pathogens in freshwater aquaculture (Hayatgheib et al., 2020). *A. hydrophila* can cause between 35-100% mortality during disease outbreaks (Baumgartner et al., 2018; Pridgeon and Klesius, 2011; Rasmussen-Ivey et al., 2016). Under experimental conditions, *A. hydrophila* can cause between 50 to 80% mortality in Nile tilapia (Abass et al., 2018; Dawood et al., 2020; Suprayudi et al., 2017). In the present study, relatively high mortality (75 −85%) was observed in immersion challenges with a MDR *A. hydrophila*. Interestingly, multiple NB-O_3_ treatments were effective with RPS of 64.7 −66.7%. The RPS value in this study was similar or higher than several studies using antibiotics for Aeromonads control in Nile tilapia e.g. RPS of 60% in orally administered with Oxytetracycline 4g/kg/feed per day (Abraham et al., 2017) or RPS 25.9 % in orally fed with Oxytetracycline 60 mg/kg biomass per day (Julinta et al., 2017).

Compared to other alternatives to antibiotics, NB-O_3_ offered comparable protective efficacy to some probiotic-based products against *Aeromonas* sp. AC9804 infection such as *Lactobacillus rhamnosus* which reported RPS values of 66.7% (Ngamkala et al., 2010) and *L. plantarum* with an RPS of 64% (Dawood et al., 2020). The results of this study were also comparable to some plant-based products used to control *A. hydrophila*, with reported RPS around 71% for Indian ginseng, *Withania somnifera* powder (Zahran et al., 2018), 35.3% for American ginseng, *Panax quinquefolius* (Abdel-Tawwab, 2012), and 58.7% for ginger, *Zingiber offcinale* (Payung et al., 2017). Our finding suggests that NB-O_3_ treatments could be considered a potential non-antibiotic approach or an “alternative to antibiotics” to control bacterial disease in aquaculture.

Ozone is among the most powerful oxidant known with oxidative potential of 2.07 volts, nearly twice of chlorine (Hugo et al., 1999). Further, aqueous ozone can generate hydroxyl radicals (OH^-^) with higher oxidative potential (2.83 volts) than ozone (Qingshi et al., 1989). Ozone ruptures cells by destroying the glycoproteins and glycolipids on the cell membranes. Moreover, ozone attacks the sulfhydryl groups of enzymes results in disruption of normal cellular enzymatic activity and loss of function. Lastly, ozone can directly damage the purine and pyrimidine bases of nucleic acids (Megahed et al., 2018). When NBs collapse, they generate shock waves that consequently lead to the formation of hydroxyl radicals (Fan et al., 2020; Takahashi et al., 2007). Thus, NB-O_3_ may enhance the disinfectant efficacy of ozone in aquaculture systems.

Although the differences in bacterial concentration in the Ah + NB-O_3_ group were only 1.0 to 1.6 fold lower than the Ah + no NB-O_3_ group after each treatment, clear differences in survivability of the fish were observed in these groups. It is also possible although not statistically significant on an individual basis the overall upregulation of innate immune genes and stimulation of humoral immune response for fish in the NB-O_3_ treatment group partially contributed to better survival rates after bacterial challenges. This has been reported by others as well (Linh et al., 2021). The stimulation of innate immunity is the first line of defense against invading pathogens and leads to improvements in health conditions and resistance to pathogens of fish (Magnadóttir, 2006). Pro-inflammatory cytokines, particularly *TNF-a* is an important macrophage-activating factor produced by leukocytes (Whyte, 2007), while lysozyme is a vital defense molecule of fish immune system due to make the demolition of bacterial cell wall (Saurabh and Sahoo, 2008). In addition, heat-shock proteins have a function in the development of specific and non-specific immune response to infections (Roberts et al., 2010).

Another factor which may also have improved survival of fish in this experiment was the DO in treated groups. Higher level of DO in NB-O_3_ treated groups during and after treatments may improve fish health by maintaining or improving normal physiological functions. Previous studies suggested that high level of oxygen improved the immunocompetence in fish (Bowden, 2008; Cecchini and Saroglia, 2002). Romano et al. (2017) revealed that 12 −13 mg/L oxygen increased immune response performance of sea bass (*Dicentrarchus labrax*).

One of the limitations of this study was our small sample size which could account for the non-significant difference in the up-regulation of innate immune genes between groups. Further, due to the limited facilities, we were unable to compare effectiveness of different forms of ozone bubbles (macro-, micro- and nanobubbles) in reducing bacterial loads and improving fish survival rate upon bacterial infection. Further studies should explore these issues to gain better understanding of this promising technology. In addition, the MRS-NB-O_3_ system need to be scaled up to be utilizable in aquaculture systems.

Despite these limitations, this study reported a MRS coupled with NB-O_3_ technology was successful at reducing mortality in fish and not exposing fish to high levels of ozone. It may be possible to scale this system up for use in hatcheries and commercial farms that use RAS systems. Our MRS-NB-O_3_ allowed multiple NB-O_3_ treatments without obvious negative impacts on the fish. This system not only suppressed MDR bacterial loads in the culture tanks, but also improved fish survivability. Application of NB-O_3_ may be a promising non-antibiotic method of reducing the risk of infectious diseases caused by bacteria, including MDR bacterial strains.

## Acknowledgement

This study was financially supported by the UK government-Department of Health and Social Care (DHSC), Global AMR Innovation fund (GAMRIF) and the International Development Research Center (IDRC), Ottawa, Canada. Le Thanh Dien received ASEAN and NON-ASEAN Scholarships of Chulalongkorn University and TSRI Fund, Chulalongkorn University_FRB640001_01_31_6. We thank to William Chalmers, City University of Hong Kong, for proofreading the manuscript.

## Disclaimers

The views expressed herein do not necessarily represent those of IDRC or its Board of Governors.

## CRediT authorship contribution statement

**Le Thanh Dien**: Conceptualization, Investigation, Methodology, Formal analysis, Writing-original draft. **Nguyen Vu Linh**: Investigation, Methodology. **Pattiya Sangpo**: Investigation. **Saengchan Senapin**: Data curation, review & editing, **Sophie St-Hilaire**: Conceptualization, review & editing, Funding acquisition, **Channarong Rodkhum**: Supervision, Validation, review & editing. **Ha Thanh Dong**: Conceptualization, Data curation, Writing-review & editing, Supervision, Validation, Funding acquisition, Project administration.

---

**Table S1.**
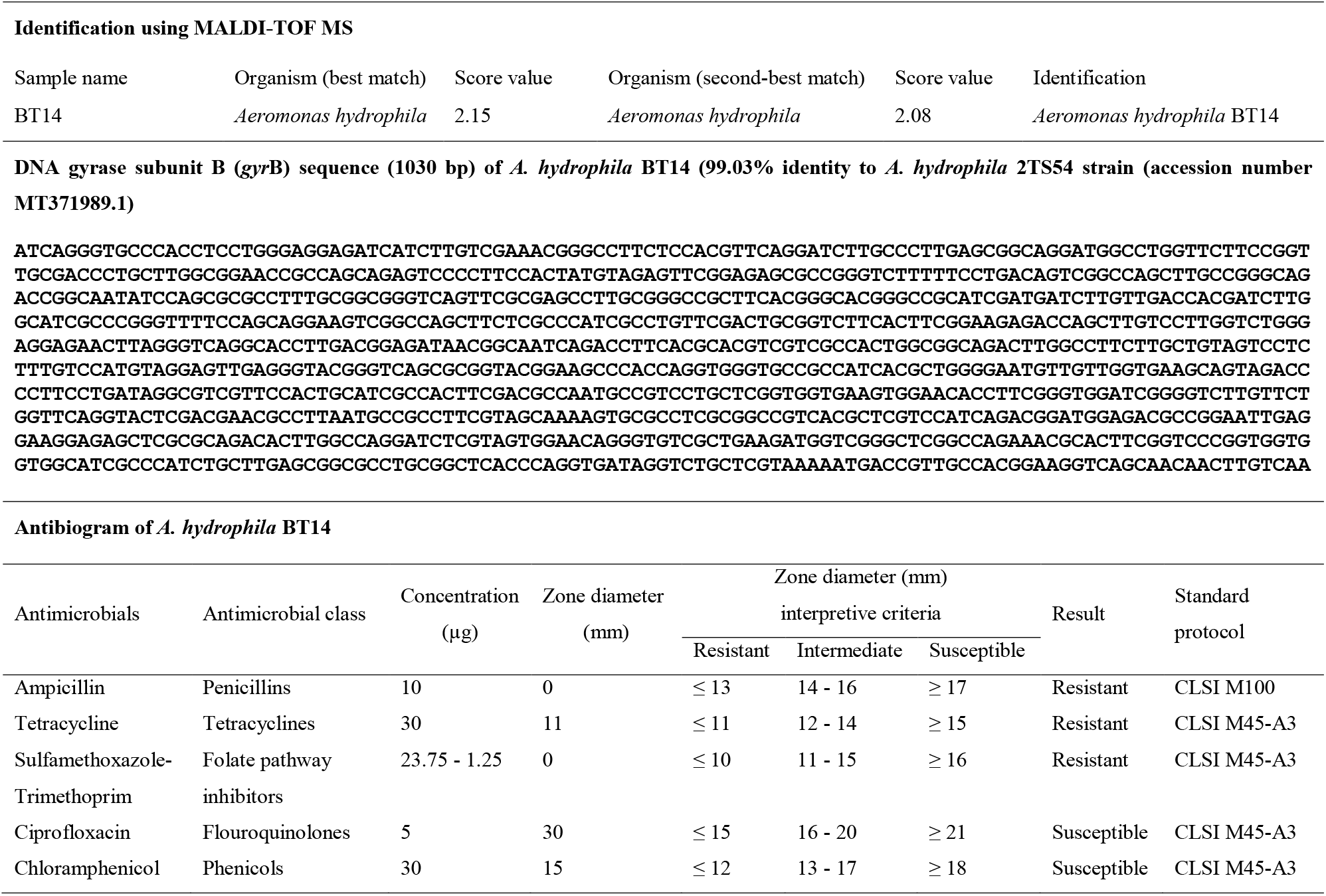
Identification and antibiogram of *A. hydrophila* BT14.

**Table S2.** Water parameters in Nile tilapia culture tank during 10 min NB-O_3_ treatment in MRS

| Treatment | Measurement time | Temperature<br>(°C) |  | pH |  | DO<br>(mg/L) |  | ORP<br>(mV) |  |
| --- | --- | --- | --- | --- | --- | --- | --- | --- | --- |
|  |  | Control | NB-O<sub>3</sub><br>treatment | Control | NB-O<sub>3</sub><br>treatment | Control | NB-O<sub>3</sub><br>treatment | Control | NB-O<sub>3</sub><br>treatment |
| 1<sup>st</sup> | Before treatment | 28.1 ± 0.2 | 27.7 ± 0.0 | 7.76 ± 0.03 | 6.64 ± 0.42 | 6.58 ± 0.08 | 6.97 ± 2.21 | 470.4 ± 3.1 | 421.3 ± 1.0 |
|  | 10 min treatment | 28.1 ± 0.3 | 28.3 ± 0.0 | 8.06 ± 0.06 | 8.03 ± 0.05 | 6.57 ± 0.04 | 15.33 ± 1.76 | 412.8 ± 0.1 | 367.8 ± 7.5 |
|  | 10 min post treatment | ND | 28.6 ± 0.0 | ND | 7.98 ± 0.03 | ND | 12.69 ± 0.68 | ND | 328.4 ± 6.1 |
| 3<sup>rd</sup> | Before treatment | 30.3 ± 0.0 | 28.6 ± 0.1 | 7.89 ± 0.03 | 7.63 ± 0.08 | 4.98 ± 0.42 | 5.82 ± 0.04 | 326.7 ± 3.8 | 465.5 ± 0.7 |
|  | 10 min treatment | 30.5 ± 0.1 | 29.2 ± 0.1 | 7.98 ± 0.08 | 7.92 ± 0.17 | 4.82 ± 0.40 | 12.29 ± 0.88 | 323.8 ± 1.8 | 387.2 ± 8.2 |
|  | 10 min post treatment | ND | 29.3 ± 0.1 | ND | 7.97 ± 0.16 | ND | 9.55 ± 1.24 | ND | 350.6 ± 11 |
| 5<sup>th</sup> | Before treatment | 29.8 ± 0.1 | 29.8 ± 0.6 | 7.59 ± 0.28 | 7.82 ± 0.00 | 4.72 ± 1.25 | 4.89 ± 0.31 | 433.8 ± 1.2 | 432.8 ± 0.8 |
|  | 10 min treatment | 29.5 ± 0.1 | 30.0 ± 0.4 | 7.98 ± 0.21 | 8.08 ± 0.04 | 5.12 ± 0.40 | 12.7 ± 0.32 | 405 ± 10.3 | 400.3 ± 9.0 |
|  | 10 min post treatment | ND | 30.3 ± 0.2 | ND | 8.14 ± 0.02 | ND | 9.28 ± 0.80 | ND | 386.2 ± 1.1 |
| 7<sup>th</sup> | Before treatment | 29.1 ± 0.1 | 29.4 ± 0.3 | 7.91 ± 0.03 | 7.95 ± 0.06 | 4.91 ± 0.12 | 5.39 ± 0.08 | 296.7 ± 9.3 | 309.9 ± 1.4 |
|  | 10 min treatment | 29.1 ± 0.1 | 29.7 ± 0.3 | 7.92 ± 0.06 | 8.01 ± 0.04 | 5.19 ± 0.30 | 12.26 ± 2.25 | 290.4 ± 4.6 | 310.3 ± 4.7 |
|  | 10 min post treatment | ND | 29.8 ± 0.4 | ND | 8.06 ± 0.04 | ND | 9.66 ± 1.70 | ND | 310.1 ± 4.9 |

DO: Dissolve Oxygen, ORP: Oxidation Reduction Potential, NB-O<sub>3</sub>: ozone-nanobubbles, ND: Not done

**Figure S1.**
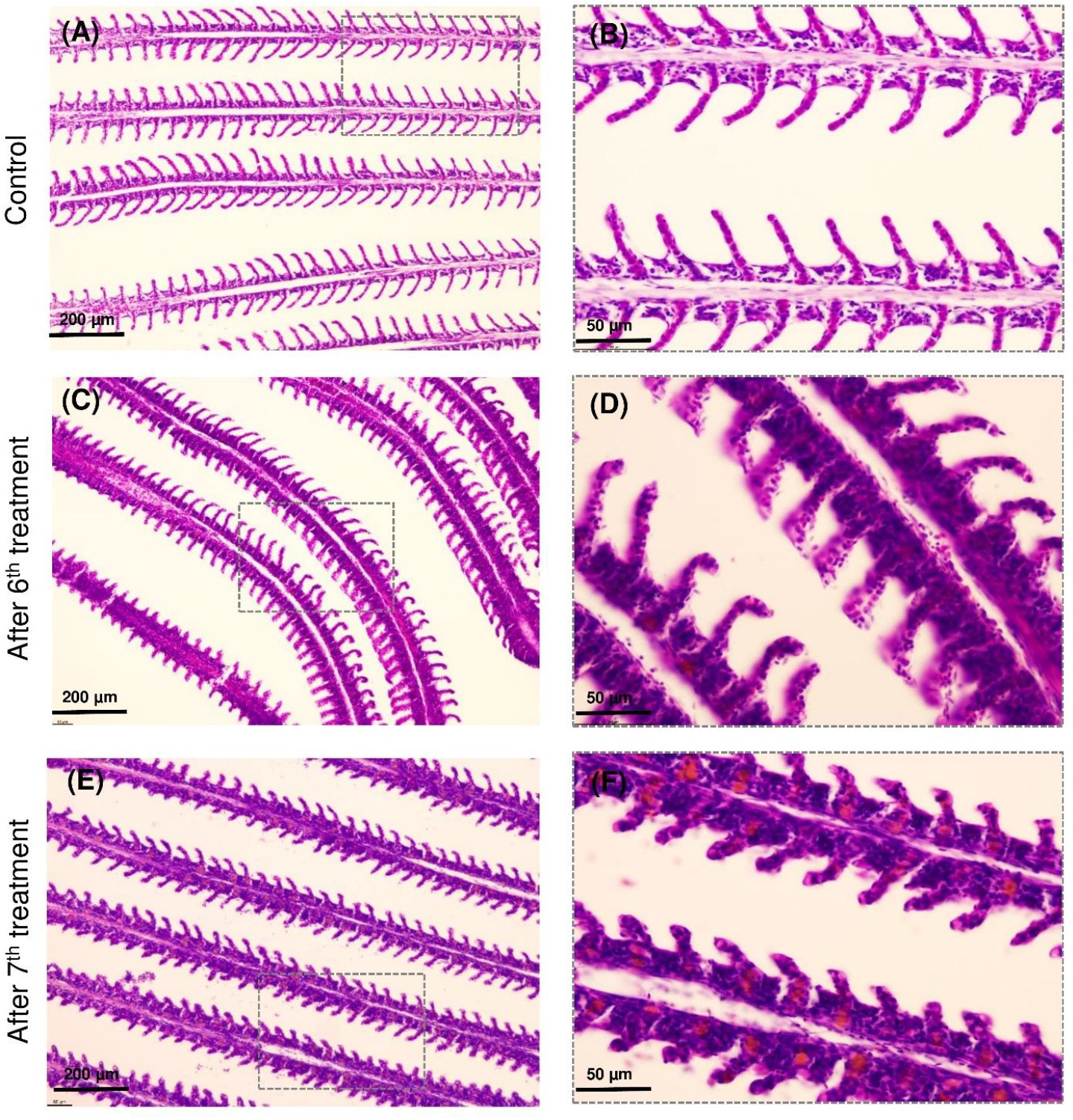
Representative photomicrographs of H&E stained sections of the gills taken at low and high magnifications. A, B, normal gill morphology from fish in control group. C, D, slight alterations in the gill lamella observed after 6^th^ treatment. E, F, alteration and increasing melanin containing cells in the gill filaments after 7^th^ treatment.

